# Holoprosencephaly and Cyclopia in *bmp7b* and *bmpr1ba* Crispant zebrafish

**DOI:** 10.1101/2024.08.09.607306

**Authors:** Valentyn Kyrychenko, Philipp Rensinghoff, Johannes Bulk, Constanze Frey, Stephan Heermann

## Abstract

Holoprosencephaly (HPE) is the most frequent developmental disorder of the forebrain. In HPE, the early single anlage of the forebrain, the Anterior Neural Plate (ANP) which encompasses the future telencephalon and eye field, fails to divide.

BMP signaling and antagonism are overall important for nervous system development. The focus of this study was on the role of the ligand *bmp7b* and the receptor *bmpr1ba* during forebrain development. The zebrafish loci of *bmp7b* and *bmpr1ba* were targeted transiently with CRISPR/Ca9. Crispants for both *bmp7b* and *bmpr1ba* presented HPE and cyclopia, one central eye. Subsequently, the ANP was addressed in *bmp7b* Crispants. The morphology of the eye field was affected, with important markers, *rx3, six3b* and *cxcr4a* expressed condensed at the midline.

Induced expression of *bmp4* is also known to result in HPE. Such *bmp4* induction altered the expression of *bmpr1ba*.

Zebrafish Crispants for *bmp7b* and *bmpr1ba* can be used as a novel HPE model. A challenge in future analyses will be the penetrance of phenotypes in Crispants. Advantages are, however, that analyses can be conducted anywhere, without the need of mutant lines. One important aspect for future analysis will be the role of individual bmp ligands, receptors and antagonists in forebrain development.

## Introduction

The development of the nervous system and especially the development of the brain is a fascinating yet intricate process. Bilateral elements of the central nervous system, e.g. the telencephalic lobes and the two retinae, are derived from single precursor domains. During neurulation, telencephalic precursors converge towards the midline, while retinal precursors stay more laterally and subsequently move even further outwards ^1^. This process depends on *rx3* and *rx3*-dependent suppression of *alcamb* (formerly *nlcam*) in retinal progenitors ^2^. It can be understood as the onset of optic vesicle out-pocketing. Forebrain development depends on BMP signaling at several stages of development. BMP antagonists are important for the induction of pre-neural identity within the future neural plate. BMP signaling activity induces epidermal fate ^3,4^. Later, during Anterior Neural Plate (ANP) development, the prospective eye field and the prospective forebrain domain compete with each other. At this stage, BMP signaling promotes telencephalic fate at the expense of eye field fates ^5^. During consecutive stages of development, BMP antagonism is essential for the proper separation of the telencephalic precursor domain, for eye field splitting and the initial out-pocketing of the optic vesicle from the forebrain ^6^. Induced expression of *bmp4* in zebrafish (at 8.5 hpf) results in anophthalmia with retinal progenitors located inside a dysmorphic forebrain, a “crypt-oculoid” ^6^. These embryos are not merely ventralized compared to embryos which received a zygotic injection of BMP ligand RNA, e.g. bmp7a/b ^7,8^. Though, early axis formation in vertebrates is strongly influenced by BMP ligands (ventral fates) and BMP antagonists (dorsal fates) organized in gradients ^9^. Knock down of BMP antagonists in zebrafish results mostly in a ventralization of the embryo ^10^. In some cases a Knock Down can, however, also result in a dorsalization of the embryo, e.g. tsg ^11^. In mouse, a double Knock Out of two BMP antagonists, Noggin and Chordin, had a pronounced but also variable effect on the morphogenesis of the head and the forebrain. Also cases of cyclopia were observed ^12^. In chicken, Holoprosencephaly (HPE) and cyclopia but not anophthalmia were reported after implantation of beads, soaked with BMP (Bmp4 and Bmp5)^13^. In zebrafish, *bmp7b* as well as specific BMP receptors (e.g. *bmpr1ba*) were transcriptionally repressed after *bmp4* induction^14^. Thus, the question was, whether a repression of e.g. *bmp7b* and of *bmpr1ba* could be involved or responsible for the phenotypes which resulted from *bmp4* induction. These phenotypes were e.g. HPE, anophthalmia, morphogenetic defects during optic cup and optic fissure formation which resulted in coloboma ^6,15–17^. In a double Knock Out for Bmpr1b and Bmpr1a in mouse, a mild form of HPE was observed, accompanied by microphthalmia ^18^. Owen and colleagues observed coloboma in zebrafish Morphants for *bmpr1ba* ^19^. Several Bmp7 mouse Knock Outs resulted in microphthalmia, anophthalmia or affected the formation of the optic fissure ^20–24^. In zebrafish, two *bmp7* paralogues were identified. The gene originally termed *bmp7* (the mutant for this is snailhouse, *snh*) is now termed *bmp7a*. In *snh* mutants a mild microphthalmia is described ^8^. The more recently identified paralog is termed *bmp7b* ^25^. Zebrafish *bmp7b* shows a closer homology to human and mouse Bmp7 ^25^. Besides the *snh* mutant, CRISPR/Cas induced mutants for both paralogs have been established recently ^26^. Notably, while *bmp7a* mutants died during gastrulation, no strong and early effect regarding eye development was observed in *bmp7b* mutants. Our group has as well targeted the *bmp7b* locus in zebrafish with the aim to generate a mutant line. Surprisingly, in homozygous in-crossed embryos, however, no obvious developmental defect was found other than a mild delay in pigmentation (unpublished). Assuming that genetic compensation ^27,28^ prevented stronger phenotypes, this approach was not followed further. Buglo and colleagues ^29^ demonstrated that genetic compensation mechanisms can be circumvented by using an acute approach by analysis of CRISPR/Cas injected zebrafish embryos (Crispants). In this study, a transient targeting strategy of the zebrafish *bmp7b* and *bmpr1ba* loci with subsequent analysis of the injected embryos (Crispants/ F0 generation) was performed. Crispants for both genes presented HPE and cyclopia. Further analysis of *bmp7b* Crispants revealed that the eye field separation was hampered. Based on these findings, a new zebrafish HPE model is presented.

## Materials and Methods

### Zebrafish care

Zebrafish were maintained at 28°C in a 12h light/ 12h dark cycle in a recirculating system. All embryos which were used for the study were offspring from zebrafish of the own facility. The following transgenic line was used: tg(hsp70l:bmp4, myl7:eGFP)^17^. Zebrafish embryos were grown in petri dishes containing zebrafish medium, 0.3g/l sea salt in deionized water.

#### Ethics statement

The permit for housing and breeding was received. Experiments in this study were performed on embryos only. These were analyzed until maximum day 3 (72 hpf). According to a directive, work with these embryonic stages do not require ethics approval.

### Heat shock procedures

For application of a heat-shock for induction of inducible transgenes, embryos were incubated in 1.5 ml reaction tubes at 37°C (heating block, Eppendorf Thermomixer). The onset of the heat shock was 8.5 hpf and the duration was 15 minutes.

### Image Processing

Images recorded by microscopy (Nikon, binocular) were edited consecutively for presentation via Inkscape (Inkscape Project. (2020). Inkscape. Retrieved from https://inkscape.org).

### In situ hybridization

Whole-mount ISHs (WMISHs) were performed according to an established protocol ^31^. Probes were partially designed cloning free ^32^. The following probes were used: emx3, foxg1a, six3b, rx3, zic2a, rx2 ^6^. bmp4 ^17^, bmp7a (F: GCAGCGGGAAATCCTCTCTA, R: GCTTCGGGACAGTTTCAGGG), bmp7b (F: GAGACGAAGAGGGCTTCTCG, R: CAGGATGACGTTGGAGCTGT), bmpr1aa (F: TAGCCAACCCCAATGCTTAC, R: TAATACGACTCACTATAGGGCCCATTTGTCTCGCAGGTAT ^33^), bmpr1ba (F: AGAATCTCTGCGGGATCTCA, R: GCTCCGTTTCTCTTGACCAG ^33^), bmpr2b (F: TATTGTCGCGCTGTTCTTTG, R: GCAGATAGGCCAGTCCTCTG ^33^), cxcr4a (F: TGGGTTGCCAGAAGAAATCC, R: CCAGAAAGGCGTACAGGATC), alcamb (F: CACCCTCTCGCTACAACTTC, R: CTTCTTGCTCTCCTCTGCTG).

### CRISPR/Cas9 F0 analysis (Crispants)

Fertilized eggs (1-cell stage embryos) were microinjected with a cocktail of Cas9 mRNA ^34^ and sgRNAs for *bmp7b* or *bmpr1ba* respectively. sgRNAs were designed using CCTop (http://crispr.cos.uni-heidelberg.de) ^35^. For *bmpr1ba* the following sgRNAs (sequences: T1: CCGTTACGCACACCTAGTGA and T2: CAGGGAACTCCAAGCAGCGG) were used with a concentration each 15 ng/µl in combination with 150 ng/µl Cas9 mRNA. For *bmp7b* the following sgRNAs (T1: CAATCCTCGGGTTACCGCAC, T4: TCTATACTCGCGATCCTCCG) were used with a concentration each 30 ng/µl in combination with 150 ng/µl Cas9 mRNA.

### Morphant analysis

A Morpholino for *bmp7b*, designed for the transition of intron 2 to exon 3 (*Bmp7b* I2:E3: TCCCTGATCGACAGACAGAGAGCGA, Gene tools, LLC) was injected into zygotes. Phenotypes were analyzed at 48 hpf. Measurements mere performed with FIJI software ^36^.

### PCR for “genotyping”

Genomic DNA was extracted from whole embryos subsequent to WMISH analysis. The following primers were used Primer Forward: ATTCACTCCGCTTGACATGTTG Primer Rev: CGCGTGACCATCTTACCTAGA. The anticipated mutant band size is 507bp. Wt and mutant band sizes are annotated in Figure 2,3 supplement.

## Results

### HPE and cyclopia in Crispants for *bmp7b* and *bmpr1ba*

The zebrafish *bmpr1ba* locus was targeted with CRISPR/Cas9. Experimental details e.g. sequences of sgRNAs are given in the Materials and Methods section. Two sgRNAs and Cas9 mRNA were co-injected into zebrafish zygotes. Phenotypes were addressed at 48 hpf. A genetically enhanced version of Cas9 was used as mRNA. This enhanced Cas9 mRNA was developed and tested rigorously ^34^. Importantly, injections of the Cas9 mRNA did not result in HPE phenotypes unspecifically. A spectrum of eye phenotypes was found in *bmpr1ba* Crsipants e.g. microphthalmia, anophthalmia, synophthalmia and importantly cyclopia (Figure 1 D, B shows a wt, Figure 1 Supplement). The penetrance of anophthalmia and cyclopia phenotypes in surviving *bmpr1ba* Crispants was approximately 6% (15/261). Less severe defects were found in approximately 5% (12/261) of surviving embryos (45% death rate). Next, the zebrafish *bmp7b* locus was targeted with CRISPR/Cas9. Experimental details e.g. sequences of sgRNAs are given in the Materials and Methods section. Two sgRNAs were injected in combination with mRNA for Cas9 into zebrafish zygotes. Phenotypes were addressed at 48 hpf (Figure 1A, scheme). The penetrance of cyclopia in surviving embryos, was approximately 14% (3/22) in *bmp7b* Crispants (58% death rate). Also, a spectrum of eye developmental defects was found, which ranged from ocular hypotelorism over synophthalmia to cyclopia (Figure 1 C, A shows a wt, Figure 1 Supplement). Occasionally, also cases of anophthalmia occurred. In Morphants for *bmp7b*, only a mild ocular hypotelorism and morphogenetic coloboma was detected (Figure 1 Supplement 2).

**Figure 1.**
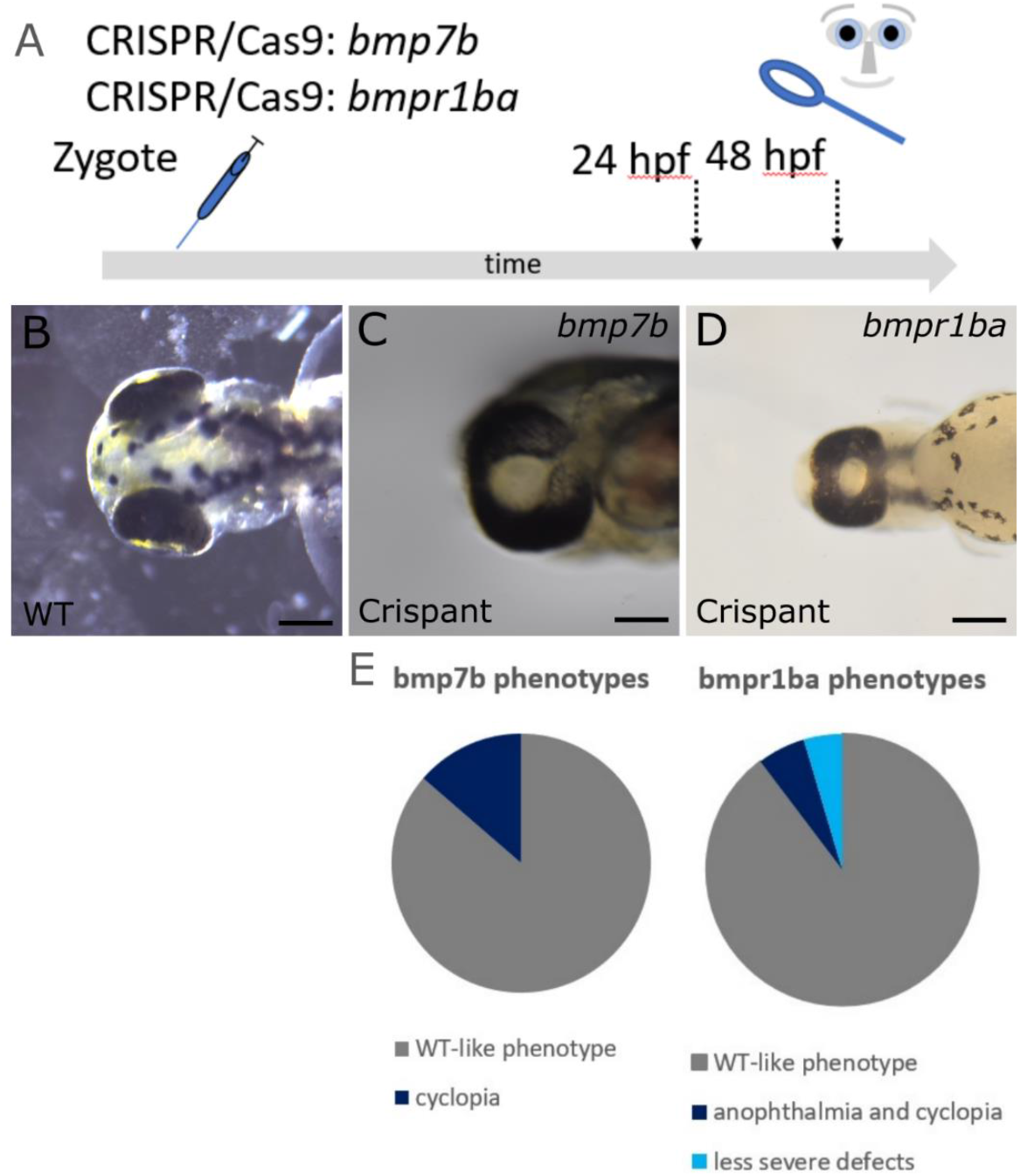
Targeting of *bmp7b* and *bmpr1ba* with CRISPR/Cas9 results in cyclopia. **A:** scheme of experimental procedure. embryos were injected at 1-cell-stage and analyzed at approx. 24-48 hpf. **B:** wildtype. dorsal view. **C:** *bmp7b* Crispant with cyclopia. ventral view. **D:** *bmpr1ba* Crispant with cyclopia. ventral view. scalebars indicate 200 µm. **E:** pie charts show the frequency of phenotypes in surviving embryos in *bmp7b* and *bmpr1ba* Crispants.

### Eye field splitting is hampered in *bmp7b* Crispants

Further analysis was performed on *bmp7b* Crispants (Figure 2, scheme of experimental design). In brief, sgRNAs and Cas9 mRNA were injected into zygotes. Subsequently, at 11 hpf, injected embryos were separated into two groups. One group of injected embryos was fixed and processed for WMISH (Figure 2, scheme of experimental design, group 1). The other group of injected embryos was grown until 48 hpf - 72 hpf. In this group the experimental efficiency of the individual experiments was estimated (Figure 2, scheme of experimental design, group 2). Un-injected embryos served as controls to record the injection-independent death rate (not shown in the scheme). The primary aim was to achieve high rates of HPE phenotypes. To this end, the dosage of CRISPR/Cas9/ sgRNA was increased. In humans, HPE is accompanied by high embryonic lethality ^37^. This suggests that in human HPE, the underlying genetic defects often result in non-viable phenotypes. Elevated levels of embryonic lethality were thus tolerated in Crispants. The experimental adaption resulted in few surviving embryos and occasionally in no surviving embryos in group 2. Death rates of individual experiments are shown in Table 1 and Table 1 supplement. The occurrence of strong eye phenotypes increased dramatically (from 14% (3/22) to 54% (91/167)). This facilitated the investigation of genes, which mark specific ANP domains (telencephalic domain and eye field) at 11 hpf (Figure 2, scheme). The main focus was the morphology of the expression domain. The secondary focus was the level of expression intensity (Figure 2, scheme of experimental design). These effects were assessed independently from each other. Binary values (“normal” and “altered”) were assigned to individual Crispant embryos, in comparison to the respective control embryos. Such binary/dichotomic analysis in which no variation was detected in the control group precluded any meaningful statistical analysis. Subsequent to this analysis, genomic DNA was extracted from the embryos and a PCR for the *bmp7b* locus was performed. A correlation between the assigned phenotype and the “Crispant genotype” was possible (Figure 2,3 supplement). A good correlation can be seen e.g. for the assigned phenotype of *six3b*, (Figure 2,3 supplement, right side, images I, J, K and L, J and L were assigned altered) and the respective genotyping results (Figure 2,3 supplement, right side, genotyping I, J, K and L, mutant bands for J and L). The experiments were carried out three times. Genes, expressed in the presumptive telencephalon, *emx3* and *foxg1a* (Figure 2 A-H, encircled domains, Figure 2,3 supplement) were found reduced in expression intensity. The morphology of the expression domain was, however, comparatively normal (Table 1, *emx3* and *foxg1a*, shaded in light grey, please see Table 1 supplement for data of individual experiments). The expression domain of genes, which mark the eye field, *six3b, rx3* and *cxcr4a* (Figure 2 I-T, encircled domains, Figure 2,3 supplement), showed an altered morphology (Table 1 and Table 1 supplement, *six3b, rx3* and *cxcr4a* shaded in dark grey). The expression domains were condensed close to the midline (*six3b*: 46,2%, *cxcr4a*: 30,8%, *rx3*: 50%) (Figure 2 J, N, R, encircled domains, Figure 2,3 supplement). This indicates that the eye field did not separate normally.

**Figure 2.**
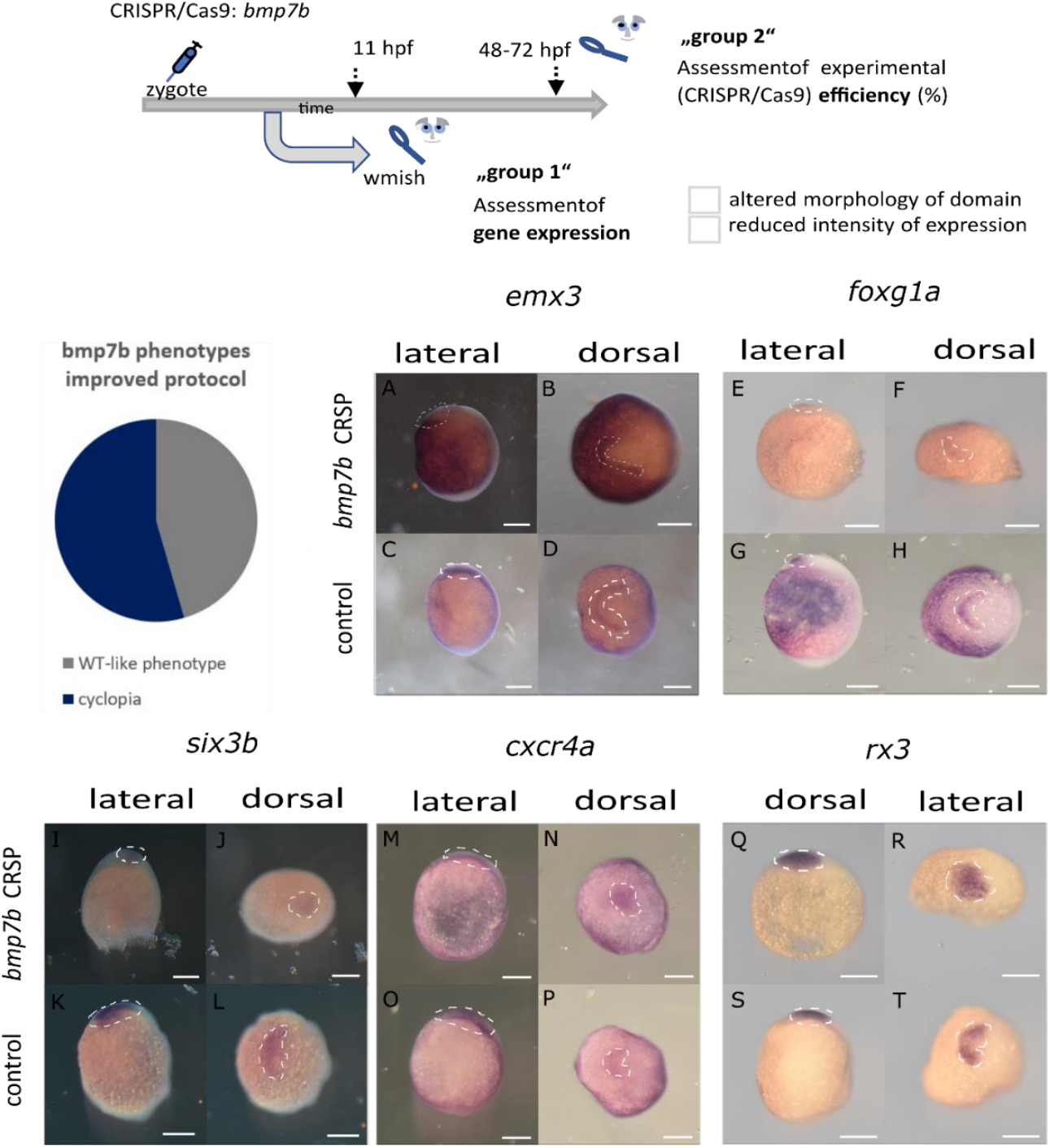
Analysis of gene expression patterns (wmish) in ANP domains in Crispants for *bmp7b* and controls (I) (embryos are presented in a lateral and in a dorsal view, the presumptive head is facing to the left, respectively, in dorsal views, expression domains are encircled). **Scheme (top) showing the experimental procedure**: CRISPR/Cas9 injections into zygotes, subsequent separation of embryos: group 1: wmish analysis, group 2: assessment of experimental efficiency (please see the **Table**) (5-6 un-injected embryos served as controls for each experiment). Three experiments were evaluated for the analysis of individual genes. Binary values were assigned to the Crispants: “normal” vs. “altered” with respect to “altered morphology of domain” and “reduced intensity of expression”, respectively. Results are given in the **Table**. Pie chart shows the frequency of phenotypes in surviving embryos with the improved protocol. **A-D** show expression patterns of *emx3* in controls (**C, D**) and Crispants **(A, B**). Please note the reduction of expression in the lateral domain (right) in **B. E-H** show expression of *foxg1a* in controls (**G, H**) and Crispants (**E, F**). Please note the reduction of expression most seen in the lateral domain (right) in **F. I-L** show expression of *six3b* in controls (**K, L**) and Crispants (**I, J**). Please note the condensed expression domain closer to the midline (**J). M-P** show expression of *cxcr4a* in controls (**O, P**) and Crispants (**M, N**,). Please note the condensed expression domain closer to the midline (**N). Q-T** show expression of *rx3* in controls (**S, T**) and Crispants (**Q, R**). Please note the condensed expression domain closer to the midline (**R)**.

**Table 1.** results of ANP analyses. Experimental design is depicted and explained in Figure 2. The respective experimental efficiency (analysis of group 2 of embryos) is given as mean of three experiments. Please note that 100% is considered either phenotype in 100 % of surviving embryos or no survivors. The data of 3 experiments (Table 1 supplement) were combined. Values are given as percentages and numbers (No).

| Gene of Interest | Experimental efficiency | No of injected embryos for analysis (wmish) (group 1) | Death-rate (group1) | Altered morphology of domain % | Altered morphology of domain No | Reduced expression intensity % | Reduced expression intensity No |
| --- | --- | --- | --- | --- | --- | --- | --- |
| *emx3* | 64% | 16 | 92,23% | 6,3 | 1 | **62,5** | **10** |
| *foxg1a* | 58% | 12 | 89,70% | 0,0 | 0 | **91,7** | **11** |
| *six3b* | 53% | 13 | 92,20% | **46,2** | **6** | 38,5 | 5 |
| *cxcr4a* | 58% | 13 | 88,03% | **30,8** | **4** | 30,8 | 4 |
| *rx3* | 81% | 6 | 89,83% | **50,0** | **3** | 0,0 | 0 |
| *alcamb* | 59% | 13 | 88,03% | 0,0 | 0 | 92,3 | 12 |
| *zic2a* | 53% | 13 | 88,47% | 46,2 | 6 | 69,2 | 9 |

The expression intensity of *alcamb* (formerly termed *nlcam*) was reduced in Crispants (Figure 3 A-D, encircled domains, Figure 2,3 supplement). The morphology of the expression domain of *zic2a*, an HPE related gene, was altered. An extended width between the lateral expression domains was observed. Additionally, a transverse domain in the posterior ANP, as seen in controls, was largely reduced in Crispants (Figure 3 E-H, encircled domains, Figure 2,3 supplement).

**Figure 3.**
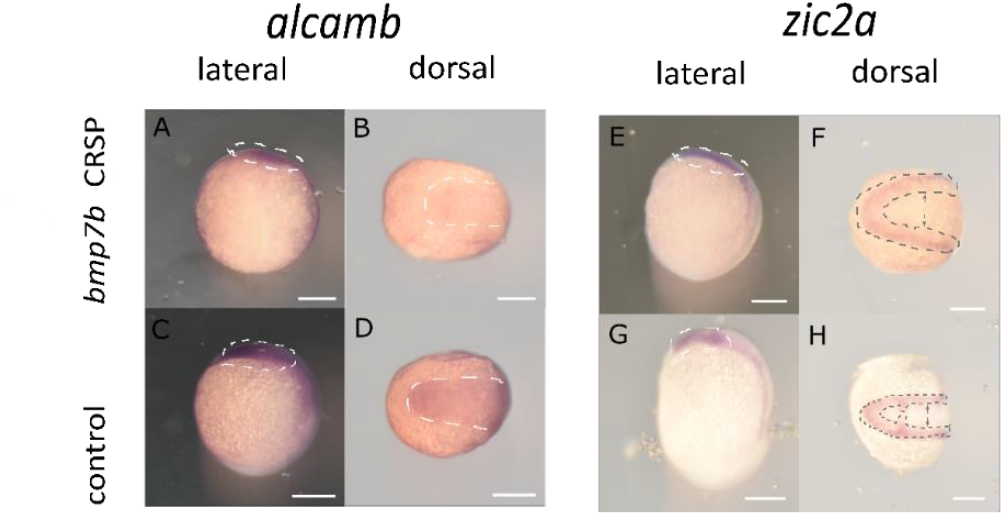
Analysis of gene expression patterns (wmish) in ANP domains in Crispants for *bmp7b* and controls (II). (embryos are presented in a lateral and in a dorsal view, the presumptive head is facing to the left, expression domains are encircled). **The experimental design is as presented in Figure 3. A-D** show expression of *alcamb* in controls (**C, D**) and Crispants (**A, B**). Please note the reduced expression in **B** in comparison to **C** (encircled). **E-H** show expression of *zic2a* in controls (**G, H**) and Crispants (**E, F**). Please note the extended width in between the lateral expression domains (arrows in **F** in comparison to arrows in **H**) as well as the reduced expression at the prospective posterior domain of the ANP (encircled in **H**, almost absent in **F**).

### Expression domain of *bmpr1ba* is affected by induced levels of *bmp4*

It was demonstrated previously that an induction of *bmp4* oversaturates endogenous BMP antagonists and results in HPE with anophthalmia ^6^. Next, the expression of *bmp7b* and *bmpr1ba* was addressed, consecutive to an induction of *bmp4*. An established transgenic line was used which expressed *bmp4* after the application of a heat-shock ^6,17^. Transgenic fish were crossed with wildtype fish. Heat-shocks were performed on embryos at 8.5 hpf. Embryos were processed for WMISH at 11 hpf (Figure 4 scheme). The expression of *cmlc2:GFP* (bleeding-heart (BH), marker for the transgenic construct) was not yet active at 11 hpf. WMISH for *bmp7b* and *bmpr1ba* was thus performed blinded. Subsequently, the embryos were genotyped. The expression intensity as well as the expression domain of *bmp7b* was not significantly altered after induction of *bmp4* (Figure 4 C, C’ and A, A’ as control). The expression domain of *bmpr1ba* was changed (Figure 4 D, D’ and B, B’ as control). In wildtype embryos (BH-negative) the expression domain was split into an anterior and a posterior domain (Figure 4 B, B’). The region in between corresponded to the prospective hypothalamic domain and showed reduced to absent levels of expression (Figure 4 B, B’). Induction of *bmp4* resulted in a continuous expression domain (Figure 4 D, D’). Overall, expression intensity seemed slightly reduced. Expression in the presumptive hypothalamic domain was enhanced.

**Figure 4.**
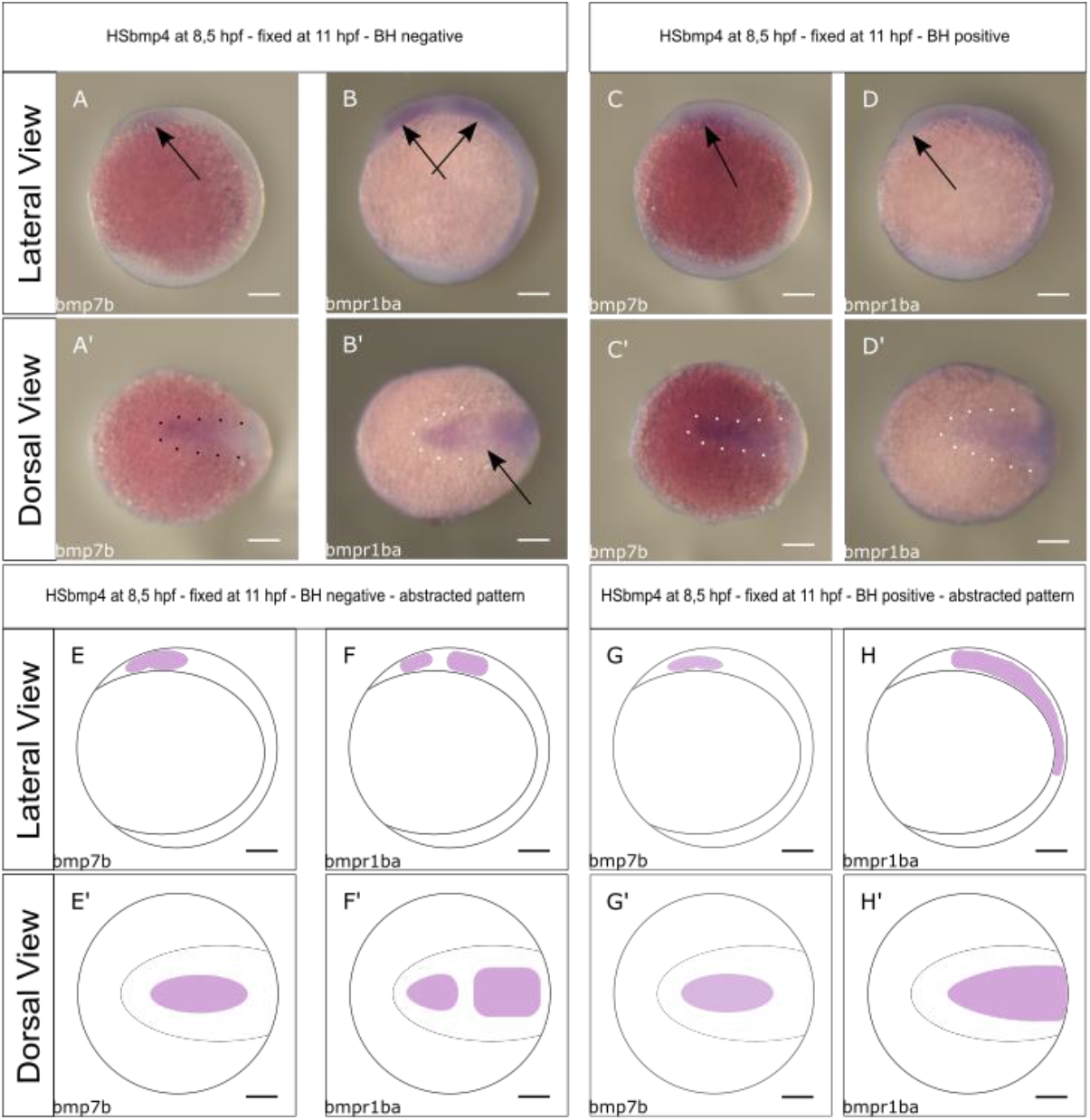
Expression patterns and domains of *bmp7b* and *bmpr1ba* in inner-embryonic domains and structures after induction of *bmp4* and in controls (lateral and dorsal views, the prospective head facing left). Scalebars indicate 200 µm. **Scheme of experimental procedure (top)**: Heat shock of outcrossed transgenes *tg(hsp:bmp4/ cmlc2:GFP)* at 8.5 hpf and fixation for whmish at 11 hpf. Genotyping and separation of recorded expression patterns **A-B’:** specimens with typical expression patterns, genotyped cmlc2:GFP/ bleeding heart (BH) negative. **C-D’:** specimens with typical expression patterns genotyped cmlc2:GFP/ bleeding heart (BH) positive. **E-H’:** schematic drawing of the respective patterns. **A, A’, E and E’:** expression pattern of *bmp7b* without *bmp4* induction (3 embryos, 3 embryos with pattern). Note the broad expression domain, at the anterior midline, likely corresponding to the anterior neuroectoderm. **B, B’, F and F’:** expression domain of the *bmpr1ba* without *bmp4* induction (5 embryos, 5 embryos with pattern). Please note the two distinct and separate expression domains. The anterior domain, likely corresponds to the anterior ANP. The posterior domain likely corresponds to the region of the midbrain-hindbrain boundary. **C, C’. G and G’:** expression domain of *bmp7b* after induction of *bmp4* (8 embryos, 6 embryos with pattern). The expression domain remains similar to the uninduced one, albeit with a slight smaller projection dorsally, seen in the lateral perspective (please note the arrow in C). **D, D’, H and H’:** expression domain of *bmpr1ba* after induction of *bmp4* (16 embryos, 16 embryos with pattern). Please note the single expression domain, which mostly includes the domains seen in the control. No gap is seen separating an anterior and a posterior expression domain as seen in the control. The anterior margin of the ANP lacks expression (arrow in D), please see B as control.

### Summary and Discussion

During early forebrain development (prosencephalon) a single central domain, which contains the presumptive telencephalon, eye field and hypothalamus, must be split, to allow the separation in two telencephalic lobes and two retinae. Pathologies during this event result in a forebrain disorder, termed holoprosencephaly (HPE). Various genetic risk factors for HPE and also some environmental risk factors have been described ^38,39^. The clinical manifestation of HPE is variable in severity and also presents a spectrum of ocular phenotypes with different severities ^40^. BMP antagonism is essential for consecutive steps during eye development, e.g. eye field splitting, optic vesicle out-pocketing ^6^ and optic vesicle to optic cup transformation^41^. Thus, BMP signaling and BMP antagonism link ocular pathologies of different severities like anophthalmia and morphogenetic coloboma, both associated with HPE. In the beforementioned analyses, BMP antagonists were oversaturated by induction of *bmp4*. In the current analysis the focus was on Crispants for *bmp7b* and *bmpr1ba*. These genes, among others, were found transcriptionally repressed after an induction of *bmp4* ^14^. HPE related eye phenotypes like cyclopia and synophthalmia were found in Crispants for both *bmp7b* and *bmpr1ba*. This indicates that both genes are essential for eye field splitting. This is well supported by the finding, that genes, that mark the eye field (rx3, cxcr4a and six3b) were found condensed at the midline in *bmp7b* Crispants (Figure 5 summary). The data suggest that *bmp7b* is acting via the BMP receptor *bmpr1ba*. An induction of *bmp4* affected the expression of *bmprb1*, which was no longer absent from the presumptive hypothalamic domain. The relevance of this observation is not yet clear. What has been shown before, is that a subduction of the presumptive hypothalamic domain plays as major role for forebrain splitting^42^.

**Figure 5.**
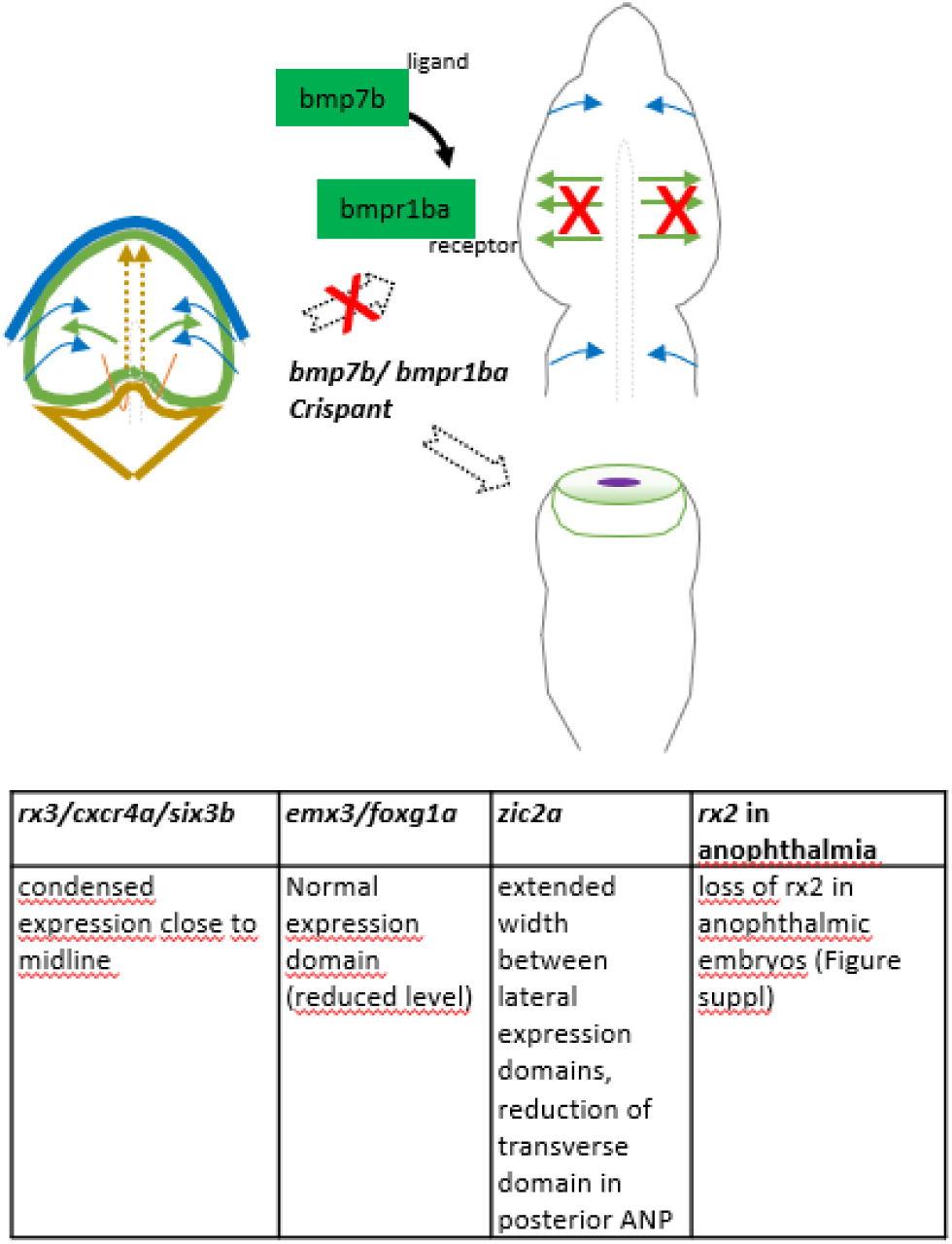
Schematic and tabular summary of major findings: In Crispants for *bmp7b* (and *bmpr1ba*) the separation of the eye field fails during early development and the development results in cyclopia. Genes expressed in the eye field *rx3, cxcr4a* and *six3b* are condensed to the midline in Crispants (shown for *bmp7b*). The morphology of the *zic2a* expression domain of was also found altered in Crispants. Zic2 was found to be important for prechordal plate development and is thus a HPE related gene ^47^. In Crispants, however, the domain was showing an extended width, while the domain marking the posterior end of the ANP was largely missing. This domain is corresponding to the presumptive hypothalamic domain, which is important for ANP splitting via a subduction movement ^42^. *Alcamb* (formerly nlcam) which in Medaka was found to be suppressed in retinal precursors ^2^, was found homogeneously expressed in the ANP of control zebrafish embryos and reduced in Crispants. Genes expressed in the prospective telencephalic domain, *emx3* and *foxg1a*, show an overall normal morphology of their expression domain (in terms of separation from the midline) but, however, reduced levels of expression intensity. In cases of Crispants resulting in anophthalmia, no “crypt-oculoid” was found.

Interestingly, in a previous attempt to address the role of *bmp7b* in a stable mutant line, generated by CRISPR/Cas9, only a delayed pigmentation could be found (not shown). This observation was at odds with other findings. Strong phenotypes were found in Bmp7 KO mice and mild HPE related phenotypes were observed in Morphants for *bmp7b* (Figure 1 Supplement 2). A mechanism that could potentially prevent stronger phenotypes in mutant lines is genetic compensation ^27,28^. Also others targeted *bmp7b* in zebrafish to establish a mutant line ^26^. The authors observed defects in eye-development, e.g. an enlarged iris and increased thickness of the retinal pigment layer ^26^. Importantly, it was shown that mRNA for *bmp7b* could still be detected in these mutants. Persisting transcription and mRNA production is part of the well described genetic compensation mechanism ^28^. This could have obscured stronger phenotypes in these mutants.

The possibility to address reverse genetics in the F0 generation ^29,43^ provided a reasonable alternative. Buglo and colleagues demonstrated that such Crispant analysis (F0) can prevent genetic compensation ^29^. Analysis of early embryonic development (e.g. 11hpf) in Crispants, however, is also challenging. Highly effective genome targeting is crucial. To a certain extent, this could be achieved with a genetically enhanced version of Cas9 mRNA, provided by colleagues ^34^. This new Cas9, importantly, did not show unspecific effects in terms of HPE phenotypes ^34^. This new version of Cas9 facilitated the analysis at 11 hpf, however, the percentages of affected Crispant embryos was still moderate (Table 1). In addition to the usage of a potent Cas9, future analyses will likely benefit also from a usage of an increased number of sgRNAs ^43,44^. The increase in efficiency in this study went along with an increase in death rates. In humans, the incidence of HPE is approximately 1/10000 in live births. The estimated incidence of HPE per conception, however, is as much as 0.4 %. This was determined in abortions in the 1960s and 1970s ^37^. Thus, high rates of embryonic lethality were tolerated during analysis. Besides the moderate percentages of strong effects on the ANP (Table 1), a variability of eye phenotypes could be observed at later embryonic stages (Figure 2,3 supplement). Both effects can be explained by the genetic mosaic Crispants. Only binary values were assigned to the Crispant embryos in terms of “normal” vs. “altered”. This binary/dichotomic analysis did not show a variation in the control group. This precluded a statistical analysis and left qualitative data. It will be important for future analyses to either cope better with genetic compensation in stable mutant lines or to further boost efficiency in Crispant embryos.

As beforementioned, the data of the current study add to the knowledge regarding forebrain and eye development. The genes *bmp4* and *bmp7b* act differentially. Induction of *bmp4* (via an oversaturation of antagonists) and Crispants for *bmp7b*, both result in HPE. A differential BMP function is not surprising per se. Different BMP ligand subfamilies are well established and members of these families are phylogenetically related, share sequence similarities and make use of specific receptor assemblies. Besides, different families also show different interaction with matrix components, due to different biochemical properties ^45^. Even though both, *bmp4* induction and *bmp7b* Crispant, result in HPE, the accompanying eye phenotypes differ. *Bmp4* induction (8.5 hpf) results in anopththalmia and a crypt-oculoid and *bmp7b* Crispants present cyclopia. An interesting difference between these two HPE models is the control of the early eye field gene *rx3. Rx3* was virtually absent after induction of *bmp4*. It was condensed at the midline in *bmp7b* Crispants. These differences might add to explain the different ocular phenotypes associated with HPE eventually. At the current stage the relevance of the observation is not clear. Induction of *bmp4* further affected the expression of *bmpr1ba* (Figure 4). An enhanced expression in the hypothalamic domain was found. In wildtype embryos this domain was largely lacking expression. It has been shown before that this region is crucial for forebrain splitting via a subduction movement ^42^. It is conceivable that the eye field splitting defect observed after *bmp4* induction ^6^ could functionally involve *bmp7* and/ or *bmpr1ba*, even though the final outcome differs and presents anophthalmia instead of cyclopia.

The zebrafish model system is used to gain insights in brain and eye development of the human being. Of course, the basics of vertebrate development are conserved, which makes zebrafish a very powerful model. Nonetheless, there are differences, one of which is the additional genome duplication in teleosts, which did not occur in human or mice. In mice for instance, one gene for Bmp7 exists, in zebrafish on the other hand, there are two. In mouse, Knock Outs for Bmp7 result in ocular phenotypes ^20,23,46^ ranging from hampered optic fissure formation to anophthalmia. In zebrafish the paralogous gene, *bmp7b*, is considered the closer homolog to the murine Bmp7. It currently remains unclear why in zebrafish *bmp7b* Crispants HPE is associated with cyclopia predominantly, while in mouse loss of Bmp7 is associated with anophthalmia.

## Conclusion

New insights regarding the role of *bmp7b* and *bmpr1ba* during eye and forebrain development are presented. A novel HPE model is shown, which is based on Crispant analysis. This model has challenges and also advantages. A major challenge is the penetrance of phenotypes in Crispants. In future, Crispant analysis will benefit from further improvement of Cas efficiency and an increase of the number of sgRNAs. One advantage of Crispant analysis is that no mutant animals need to be raised. Thus, analysis can be conducted and reproduced anywhere, without the need to import and use mutant lines. One important aspect for future analysis will be to further dissect the role of individual BMP ligands and receptors in forebrain development especially against the background of different eye phenotypes in HPE.

## Supplementary Materials

Figure 1 Supplement

Figure 1 Supplement 2

Figure 2,3 supplement

Table 1 Supplement

**Figure 1 Supplement:**
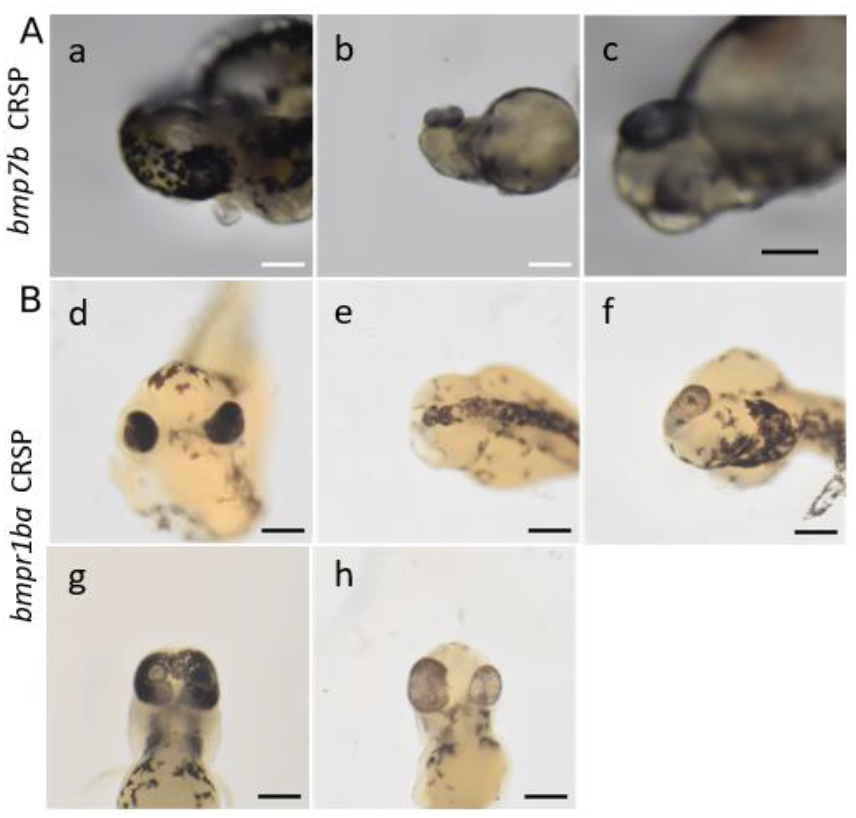
Additional phenotypes observed after targeting *bmp7b* and *bmpr1ba* with CRISPR/Cas9. **A:** *bmp7b* Crispants. **a:** cyclopia (anterior view). **b:** one lateral eye (dorsal view). **c**: synophthalmia (ventral view) **B**: *bmpr1ba*-Crispants.**d:** microphthalmia. ventral view. **e**: anophthalmia. dorsal view. **f:** one lateral eye. dorsal view. **g:** synophthalmia/cyclopia, ventral view. **h:** ocular hypotelorism. ventral view. Scalebars indicate 200 µm.

**Figure 1 Supplement 2:**
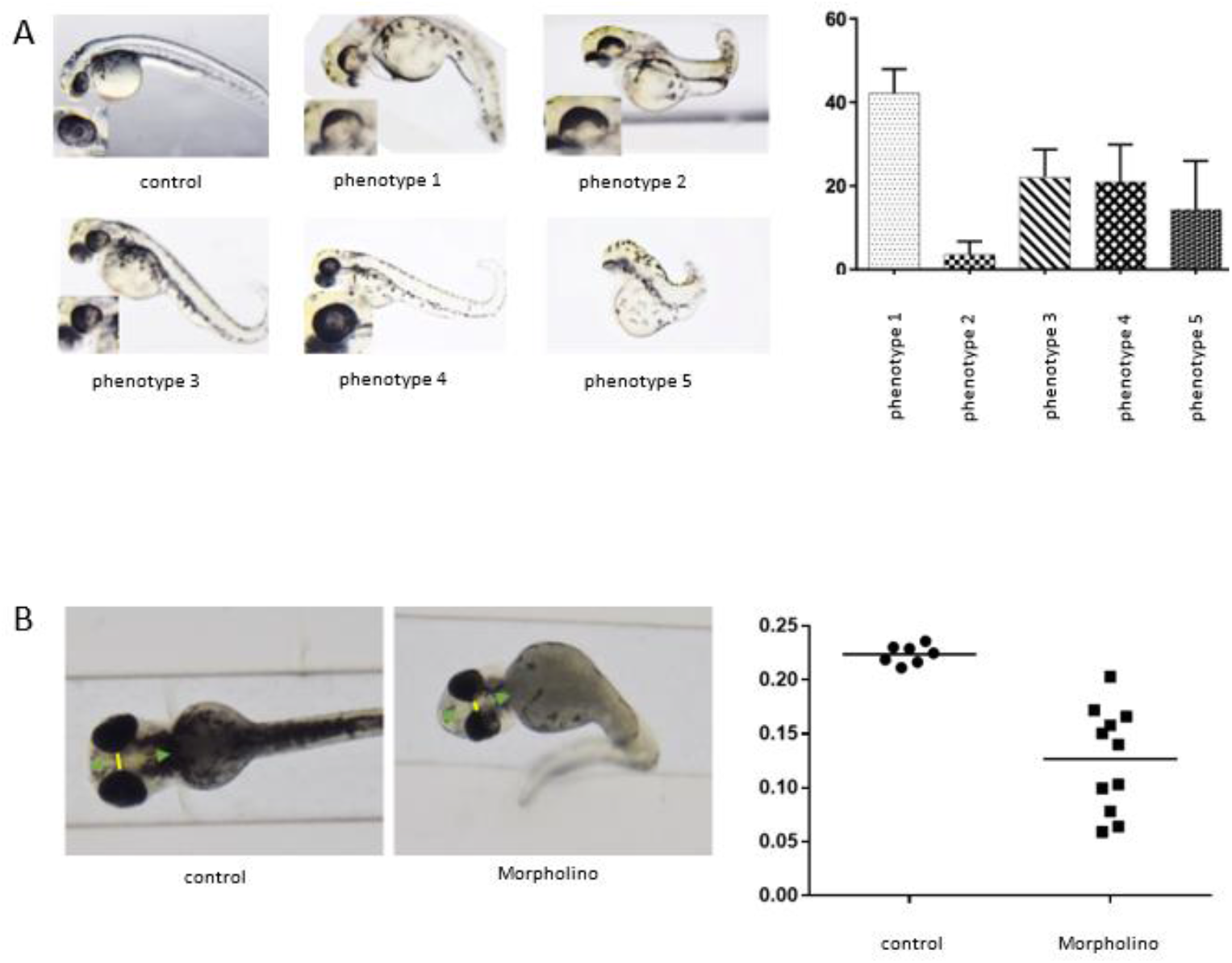
*Bmp7b* Morphant phenotypes. **A:** Control embryo and *bmp7b* Morphants (lateral view, head left, 48 hpf), optic cup formation is affected with variable intensity (phenotypes 1 to phenotype 4, close ups are presented in inserts), a broad coloboma is visible (phenotype 1, 2, 5), graph shows percentage distribution of phenotypes. **B:** control and *bmp7b* Morphant (ventral view, head left), distance between the eyes (yellow marking) was measured in relation to the length of the head (green arrow), graph shows the results as a scatter plot in µm.

**Figure 2,3 supplement:**
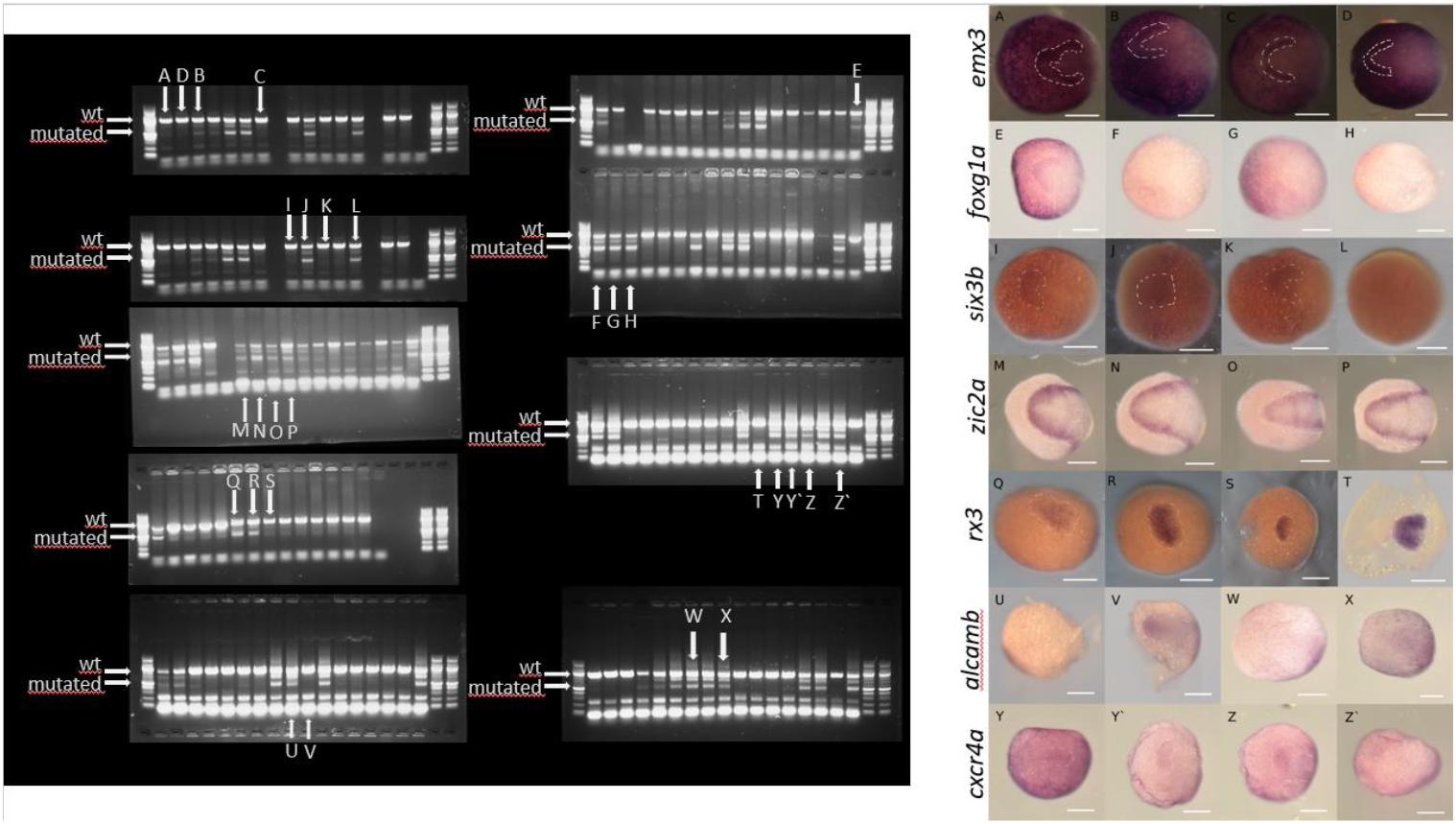
**Left side** is showing PCR analysis for the *bmp7b* genomic locus. The size of the wt band ist indicated, as well was the size of reduced mutated bands. **A-Z’** mark the PCR results for embryos, which are shown on the right side. **Right side** is showing wmish analysis of individual Crispant embryos, expression domains are encircled, information is given regarding the binary assignment („normal” vs. „altered”) **A-D**: ***emx3*** (B is considered altered, A, C, D are considered normal), **E-H**: ***foxg1a*** (F, G, H are considered altered, E is considered normal), **I-L**: ***six3b*** (J, L are considered altered, I, K are considered normal), **M-P**: ***zic2a*** (: N, P are considered altered, M,O are considered normal), **Q-T**: ***rx3*** (Q, R are considered altered, S, T are considered normal), **U-X**: ***alcamb*** (U, W, X are considered altered, V is considered normal), **Y-Z’**: ***cxcr4a*** (Y, Y`, Z` is considered altered, Z is considered normal)

**Table 1 supplement.** results of ANP analyses. Experimental design is depicted and explained in Figure 2. The respective experimental efficiency (analysis of group 2 of embryos) is given. Please note that 100% is considered either phenotype in 100 % of surviving embryos or no survivors. Data are presented per individual experiment (left) but also per gene (right). In the latter the data of the 3 experiments were combined. In addition, the respective values are given as percentages and absolute numbers (No).

| Gene of Interest | Analysis "per experiment" |  |  |  |  |  |  |  |
| --- | --- | --- | --- | --- | --- | --- | --- | --- |
|  | Experiment | Experimental efficiency (group 2) | No of injected embryos for analysis (wmish) (group 1) | Death-rate (group 1) | Altered morphology of domain % | Altered morphology of domain No | Reduced expression intensity % | Reduced expression intensity No |
| *emx3* | 1 | 25% | 7 | 89,40% | 14,30% | 1 | 28,70% | 2 |
|  | 2 | 100% | 4 | 98,50% | 0 | 0 | 100% | 4 |
|  | 3 | 66,60% | 5 | 88,80% | 0 | 0 | 80% | 4 |
| *foxg1a* | 1 | 40% | 5 | 85,30% | 0 | 0 | 80% | 4 |
|  | 2 | 58,30% | 3 | 88,30% | 0 | 0 | 100% | 3 |
|  | 3 | 75% | 4 | 95,50% | 0 | 0 | 100% | 4 |
| *six3b* | 1 | 25% | 5 | 89,40% | 40% | 2 | 20% | 1 |
|  | 2 | 100% | 4 | 98,50% | 0 | 0 | 100% | 4 |
|  | 3 | 33,30% | 4 | 88,70% | 100% | 4 | 0% | 0 |
| *cxcr4a* | 1 | 40% | 3 | 85,30% | 33,30% | 1 | 0% | 0 |
|  | 2 | 58% | 5 | 83,30% | 20% | 1 | 60% | 3 |
|  | 3 | 75% | 5 | 95,50% | 40% | 2 | 20% | 1 |
| *rx3* | 1 | 100% | 1 | 98,50% | 0% | 0 | 0% | 0 |
|  | 2 | 66,60% | 3 | 85,70% | 67% | 2 | 0% | 0 |
|  | 3 | 77% | 2 | 85,30% | 50% | 1 | 0% | 0 |
| *alcamb* | 1 | 44% | 4 | 85,30% | 0% | 0 | 100% | 4 |
|  | 2 | 58,30% | 4 | 83,30% | 0% | 0 | 75% | 3 |
|  | 3 | 75% | 5 | 95,50% | 0% | 0 | 100% | 5 |
| *zic2a* | 1 | 40% | 5 | 85,30% | 40% | 2 | 80% | 4 |
|  | 2 | 58,30% | 3 | 88,30% | 33% | 1 | 67% | 2 |
|  | 3 | 60% | 5 | 91,80% | 60% | 3 | 60% | 3 |

